# No evidence for aberrant expression of amyloid β and phosphorylated tau proteins in herpes simplex virus-infected human neurons *in vivo*

**DOI:** 10.1101/2021.09.24.461630

**Authors:** Diana N. Tran, Amy T.C.M. Bakx, Vera van Dis, Eleonora Aronica, Robert M. Verdijk, Werner J.D. Ouwendijk

## Abstract

Increasing evidence implicates the neurotropic herpes simplex virus 1 (HSV-1) in the pathogenesis of Alzheimer’s Disease (AD). However, it is unclear whether previously reported findings in HSV-1 cell culture and animal models can be translated to humans. Here, we analyzed clinical specimens from latently HSV-1 infected individuals and individuals with lytic HSV infection of the brain (herpes simplex encephalitis; HSE). Latent HSV-1 DNA load in trigeminal ganglia was identical between AD patients and controls, and latently HSV-infected neurons did not express amyloid β (Aβ) or hyperphosphorylated tau (pTau). Some HSE patient brains presented with ageing-related intraneuronal Aβ accumulations, neurofibrillary tangles (NFT) or extracellular Aβ plaques, but these were neither restricted to HSV-infected neurons nor brain regions containing virus-infected cells. Analysis of unique brain material from an AD patient with concurrent HSE showed that HSV-infected cells frequently localized close to Aβ plaques and NFT, but did not exacerbate AD-related pathology in relation to HSV infection. HSE-associated neuroinflammation was not associated with specific Aβ or pTau phenotypes. Collectively, the data indicate that neither latent nor lytic HSV infection of human neurons *in vivo* is directly associated with aberrant Aβ or pTau protein expression.

## BACKGROUND

Alzheimer’s disease (AD) is a progressive neurodegenerative disease that accounts for about 80% of dementia cases (1). Pathological hallmarks of AD are the formation of extracellular amyloid β (Aβ) protein plaques and intracellular neurofibrillary tangles (NFT) composed of hyperphosphorylated tau (pTau) protein (2). The deposition of Aβ plaques is an early event in AD pathogenesis, which induces subsequent inflammation, accumulation of NFTs, and ultimately, neuronal dysfunction and cell death (3). Genetic factors largely explain the risks of AD development and appear to be principally involved in Aβ processing and clearance – most notably apolipoprotein ε4 allele (*APOE4*) – and are expressed by CNS-resident immune cells (4). However, an estimated 21 – 42% of lifetime risk on AD is attributed to environmental factors (5). Recent studies demonstrated that Aβ can function as an antimicrobial peptide that protects neurons from bacterial and viral infections (6,7), increasing the evidence that microbial infections – especially herpes simplex virus 1 (HSV-1) – could play a role in the pathogenesis of AD.

Most adults worldwide are infected with the human neurotropic alphaherpesvirus herpes simplex virus 1 (HSV-1). Latent HSV-1 predominantly resides in the somata of pseudounipolar sensory trigeminal ganglion (TG) neurons, which innervate both the orofacial epithelia and the brainstem (8). Periodic HSV-1 reactivation results in virus spread to the oral mucosa and the central nervous system (CNS). Whereas oral HSV-1 shedding can be both asymptomatic and symptomatic (cold sores), symptomatic HSV-1 infection of the CNS is extremely rare and associated with severe morbidity and mortality (herpes simplex encephalitis; HSE) (9,10). HSV-1 DNA is detected in brain of elderly individuals, with a significantly higher frequency in brain of AD patients in which viral DNA was found to co-localize with Aβ plaques (11,12). HSV-1 infection of neuronal cell cultures induces the accumulation of intracellular Aβ and extracellular Aβ aggregates, although variable results were obtained using three-dimensional neuron models (13–17). Additionally, HSV-1 infection induces nuclear accumulation and hyperphosphorylation of tau protein in neurons *in vitro* (18–20). These AD-related pathological changes were recently found to progressively accumulate with repeated HSV-1 infection and to correlate with cognitive impairment in mice (21); however, humans are the only natural host of HSV-1 and, unlike mice, lytic HSV-1 infection of the CNS is extremely rare, at least in part because human cortical neurons are intrinsically resistant to HSV-1 infection (22). It remains to be determined whether lytic or latent HSV-1 infection of human neurons *in vivo* is directly associated with aberrant Aβ and pTau expression. Therefore, we investigated the relationship between AD-related neuropathological changes and HSV infection in TG of latently HSV-1 infected individuals and brain of individuals with lytic HSV infection of the brain (HSE).

## METHODS

### Human clinical specimens

Temporal cortex biopsies from AD patients, and paired human TG and plasma samples from healthy controls, AD patients, and patients with other neurological diseases were obtained from the Netherlands Brain Bank (Netherlands Institute for Neuroscience; Amsterdam, the Netherlands). All donors had provided written informed consent for brain autopsy and the use of material and clinical information for research purposes. All study procedures were performed in compliance with relevant laws in the Netherlands. Institutional guidelines approved by the local ethical committee (VU University Medical Center, Amsterdam, project number 2009/148) and in accordance with the ethical standards of the Declaration of Helsinki. AD brain tissue and part of TG biopsies were formalin-fixed and paraffin-embedded (FFPE) for *in situ* analysis. Additional TG samples were used to generate single cell suspensions (23). FFPE brain samples from patients with traumatic brain injury (TBI), HSE patients and a VZV encephalitis patient were obtained for diagnostic purposes and provided by the BioBanks of the Erasmus MC (TBI patients) and Amsterdam University Medical Center (HSE and VZV encephalitis). According to the institutional “Opt-Out” system, which is defined by the National “Code of Good Conduct” [Dutch: Code Goed Gebruik, May 2011], these surplus human brain tissues were available for the current study.

### DNA extraction, quantitative TaqMan Real-Time PCR (qPCR) and *APOE* genotyping

One-tenth of a TG single cell suspension was used for DNA extraction using the QIAamp DNA Kit (Qiagen). TaqMan qPCR was performed in duplicate on the 7500 Real-Time PCR system using 2X PCR Universal Master Mix (Applied Biosystems) and primer/probe pairs specific for HSV-1 *US4*, VZV *ORF38* and the human single copy gene hydroxymethylbilane synthase (24) (Supplementary Table 1). *APOE* genotyping was performed using allele ε2-, ε3- and ε4-specific primer/probe combinations and TaqMan qPCR, as described (25) (Supplementary Table 1). Results were confirmed by PCR amplification of the *APOE* gene region containing allele-differentiating SNPs rs429358 and rs7412 using PfuUltra II Fusion High-Fidelity DNA Polymerase (Agilent), followed by Sanger sequencing using the BigDye 3.1 Cycle Sequencing Kit on the 3130XL Genetic Analyzer (Applied Biosystems) (25).

### Immunohistochemical and immunofluorescent staining

FFPE tissue sections were subjected to heat-induced antigen retrieval using citrate buffer or Trilogy™ (Cell Marque). Immunohistochemical (IHC) staining was performed using the following primary antibodies: monoclonal mouse anti-β-Amyloid (clone 4G8, BioLegend), anti-phosphorylated-Tau (Ser^202^/Thr^205^) (AT8, Invitrogen), and anti-HSV-1 infected cell protein 8 (ICP8) (10A3, Cell Marque). Staining was visualized using biotinylated polyclonal rabbit anti-mouse Ig (Dako) and goat anti-rabbit Ig (Dako), followed horseradish peroxidase-conjugated streptavidin (Dako) and 3-Amino-9-ethylcarbazole substrate. Images were obtained with an Olympus BX51 microscope or by scanning the slides using the Hamamatsu NanoZoomer 2.0 HT.

Immunofluorescent (IF) staining was performed using the following primary antibodies: polyclonal rabbit anti-HSV-1 (cross-reactive with HSV-2; Agilent), anti-Iba1 (Wako) or anti-GFAP (Dako), polyclonal chicken anti-GFAP (Abcam), monoclonal mouse anti-NeuN IgG1 (A60, Sigma-Aldrich), anti-MBP (1.B.645, Santa Cruz Biotechnology), anti-Iba1 (GT10312, Invitrogen), anti-β-Amyloid (4G8, BioLegend), anti-phosphorylated-Tau (Ser^202^/Thr^205^) (AT8, Invitrogen), anti-CD45 (2B11 +PD7/26, Dako), and monoclonal rat anti-CD3 (CD3-12, Abcam). Alexa Fluor® 488 (AF488)-, AF594- and AF647-conjugated polyclonal goat anti-rabbit IgG, anti-mouse IgG1 and IgG2b, goat anti-chicken IgY and goat anti-rat IgG secondary antibodies (Invitrogen) were used. Nuclei were stained with Hoechst 33342 Solution and mounted with ProLong™ Diamond Antifade Mountant (Thermo Fisher Scientific). Images were obtained using a Zeiss LSM700 confocal microscope.

### *In situ* hybridization

Tissue sections were stained for HSV-1 and VZV RNA by *in situ* hybridization (ISH), using the RNAscope®2.5 HD Kit-RED and probes HSV-1-LAT (Cat No. 315651) and VZV-Pool (Cat No. 400701) (Advanced Cell Diagnostics). Slides were counterstained with hematoxylin (Sigma) and mounted with EcoMount (Biocare Medical).

### Statistical analyses

All statistical analyses were performed using GraphPad Prism 8.0.2 (GraphPad Software Inc).

## RESULTS

### HSV-1 DNA load in TG of AD patient and controls

Previous studies suggest that HSV-1 infection or reactivation, as measured by plasma HSV-1 IgG and IgM levels, is a risk factor for AD development (26). Moreover, clinical HSV-1 reactivation frequency correlates with both *APOE4* carriage (humans) and latent viral DNA load (mice) (27–30). To investigate whether latent HSV-1 DNA load was associated with AD development or *APOE* genotype, we performed qPCR on human TG that were infected with HSV-1 and obtained from AD patients and controls. Additionally, we performed qPCR for the closely related varicella-zoster virus (VZV), because most human TG are co-infected by HSV-1 and VZV (10). HSV-1 and VZV DNA loads were similar between AD patients and controls (Figure 1A). Further, *APOE* genotyping of all analyzed TG specimens demonstrated comparable HSV-1 and VZV DNA levels in *APOE4* carriers and non-carriers (Figure 1B and Supplementary Figure 1), despite the reported effect of *APOE4* on promoting HSV neurovirulence (32,33).

**Figure 1.**
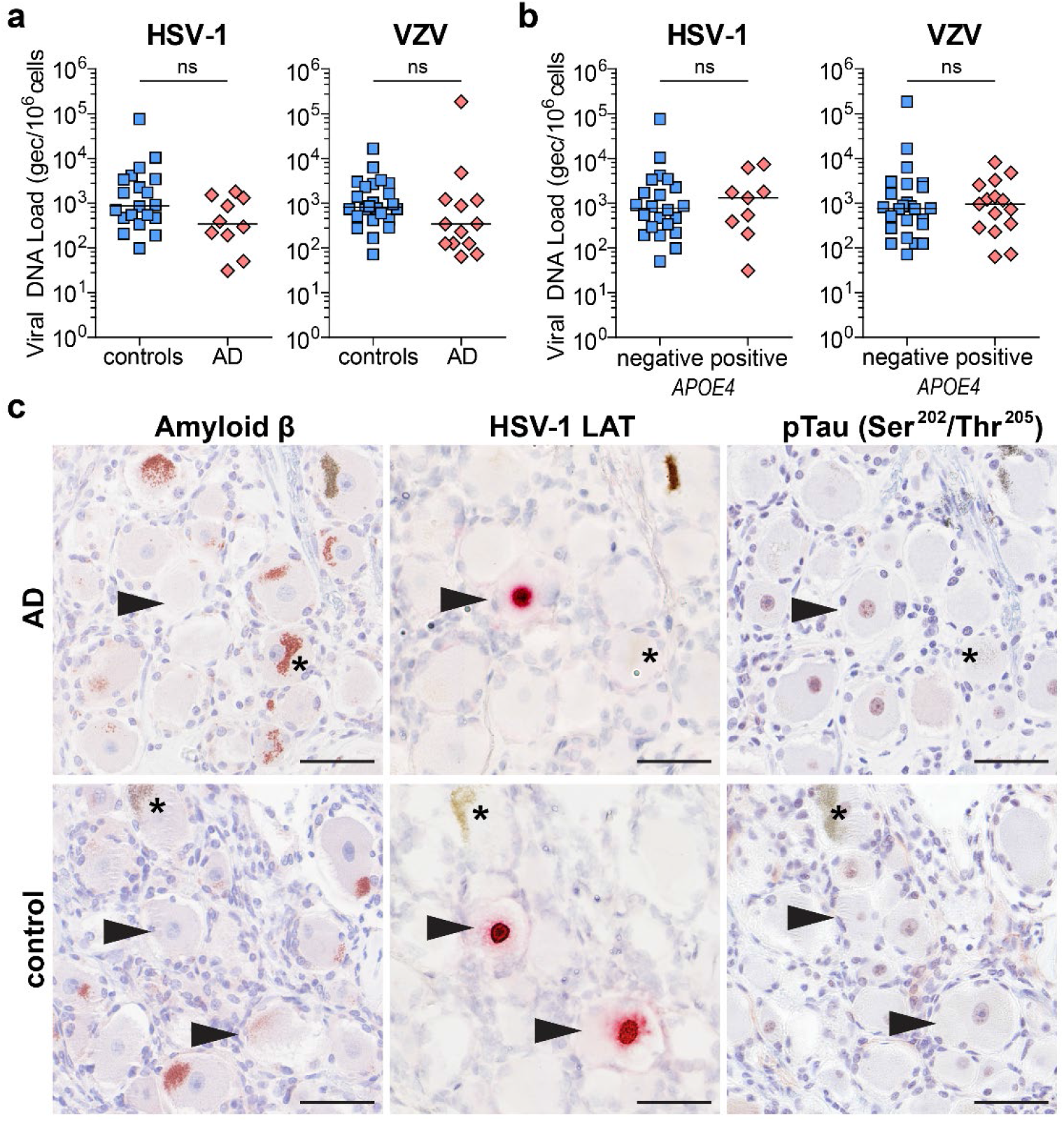
HSV-1 infection is not associated with aberrant Aβ or pTau expression in latently infected human trigeminal ganglia (TG) neurons. (**A, B**) HSV-1- and VZV-specific qPCR was performed on DNA extracted from the TG of Alzheimer’s disease (AD) patients and control subjects, stratified on disease status (**A**; 20 controls and 10 AD patients) or *APOE4* allele carrier status (**B**; HSV-1: 22 *APOE4*-negative and 8 *APOE4*-positive individuals; VZV: 24 *APOE4*-negative and 13 *APOE4*-positive individuals). Horizontal line: median. (**C**) Sequential TG sections from AD (n = 4) and control (n = 6) subjects were stained for amyloid β protein, HSV-1 latency-associated transcript (LAT) RNA and phosphorylated Tau protein (pTau; Ser^202^/Thr^205^) by immunohistochemistry (IHC) and RNA *in situ* hybridization (ISH). Arrowheads indicate LAT-positive neurons and asterisks indicate lipofuscin granules. Scale bar: 50 μm.

### No expression of Aβ and pTau in latently HSV-infected TG neurons of AD patients and controls

Most individuals acquire HSV-1 infection in their childhood, followed by a lifelong latent infection and frequent (asymptomatic) virus reactivation (34,35). Given that HSV-1 replication leads to the accumulation of Aβ and increased pTau expression in cultured neurons and in the brains of HSV-infected mice (13,14,18,21,36), we analyzed whether latent HSV-1 infection was associated with Aβ or pTau expression in TG neurons of AD patients and controls (Supplementary Table 2). Analysis of consecutive TG sections from HSV IgG seropositive (n = 3 AD; n = 5 controls) and HSV IgG seronegative (n = 2 AD) donors did not demonstrate Aβ, Aβ plaques or pTau staining in latently HSV-1-infected TG neurons of either AD patients or controls (Figure 1C). Similarly, we did not observe aberrant Aβ or pTau staining in neurons not infected by HSV-1.

### Expression of Aβ and pTau in brain of HSE patients

We hypothesized that HSV-induced AD-related pathology could be restricted to CNS neurons, rather than peripheral TG neurons, or could be reversible and therefore limited to lytic (productive) virus infection. To test these hypotheses, we acquired rare post-mortem brain specimens from five HSE patients (Table 1), encompassing 4 cases of encephalitis caused by HSV-1 and 1 neonatal HSV-2 patient. Brain sections from all patients analyzed contained lytically HSV-infected cells, as evidenced by the detection of intranuclear eosinophilic (Cowdry type A) inclusion bodies, HSV-1 LAT RNA (in all HSV-1 HSE patients) and HSV ICP8 protein (Figure 2A). In all patients, the majority of HSV-infected ICP8^POS^ cells were identified as NeuN^POS^ neurons, with lower frequencies of infected non-neuronal cells, mainly Iba1^POS^ microglia (Figure 2B-C). Thus, brain sections from HSE patients provide a snapshot of lytically HSV-infected CNS neurons in vivo.

**Table 1.** HSE patient characteristics and Alzheimer’s disease-related pathology

| Patient | Age<sup>a</sup> | Gender<sup>b</sup> | Causative<br>Virus | Brain regions analyzed | Aβ<br>intraneuronal<sup>c</sup> | Aβ<br>plaques | NFT<sup>d</sup> |
| --- | --- | --- | --- | --- | --- | --- | --- |
| 1 | -- | -- | HSV-1 | Cortex | No | No | No |
| 2 | 79 | F | HSV-1 | Hippocampus / Cortex | Yes | Yes | Yes |
|  |  |  |  | Amygdala / Entorhinal<br>Cortex | Yes | Yes | Yes |
|  |  |  |  | Cortex | Yes | Yes | Yes |
|  |  |  |  | Insula | Yes | Yes | Yes |
| 3 | 0<sup>e</sup> | F | HSV-2 | Cortex | No | No | No |
| 4 | 81 | F | HSV-1 | Putamen | No | Yes | No |
| 5 | 76 | M | HSV-1 | Hippocampus /<br>Entorhinal Cortex | No | Yes | Yes |

<sup>a</sup> Age in years; <sup>b</sup> F, female; M, male; <sup>c</sup> intraneuronal Aβ accumulation; <sup>d</sup> NFT, neurofibrillary tangle; <sup>e</sup> neonatal HSE patient, 2 weeks old

**Figure 2.**
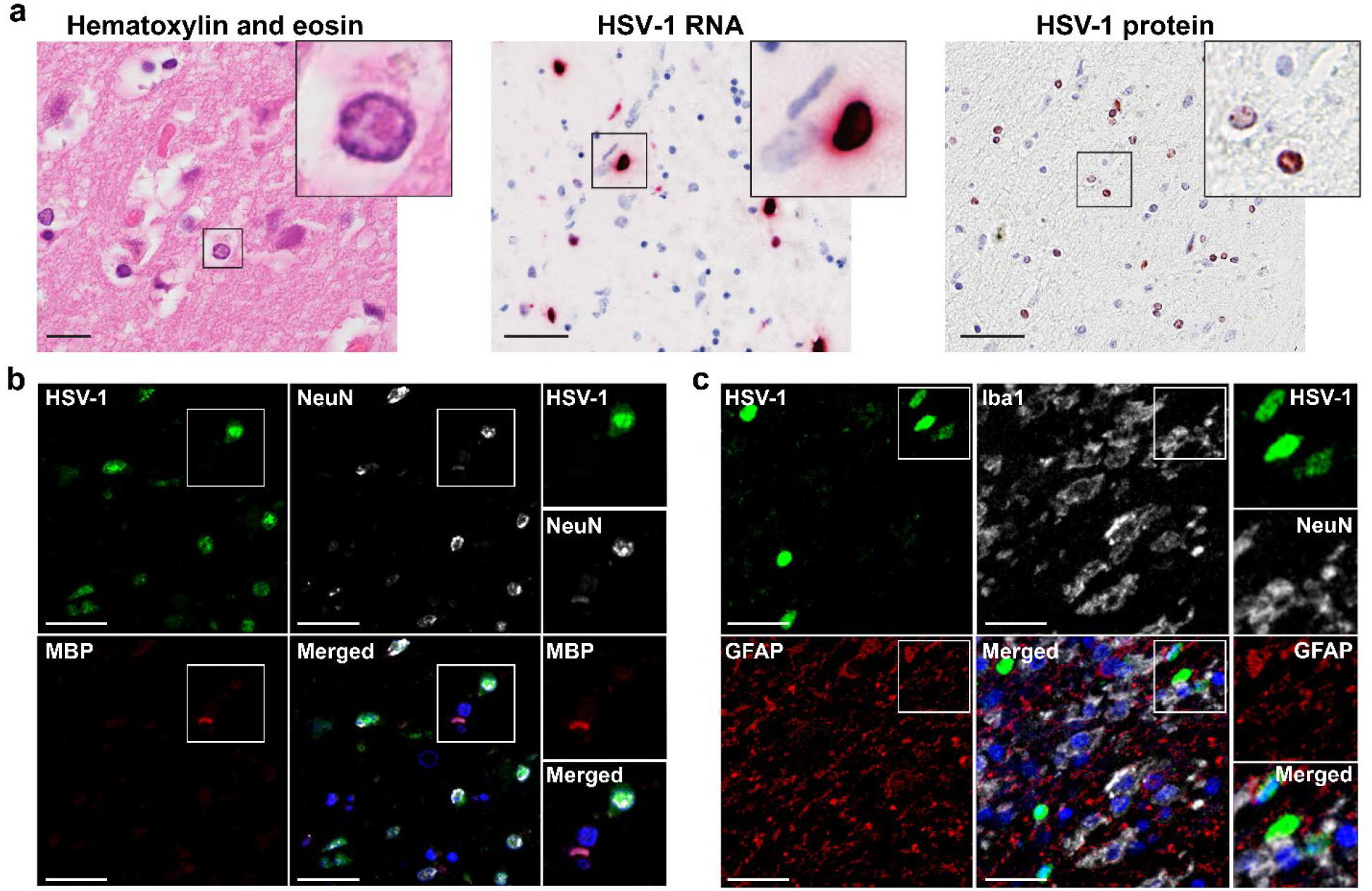
Lytic HSV infection of neurons in brain of herpes simplex virus encephalitis (HSE) patients. (**A**) Brain sections from HSE patients (Table 1) were stained with hematoxylin and eosin, or stained for HSV-1 LAT RNA by ISH and HSV-1 ICP8 protein by IHC. Boxes indicate area shown at higher magnification in the inset. Scale bar: 50 μm. (**B, C**) Brain sections of HSE patients were immunofluorescently stained for (**B**) HSV-1 protein (green), neurons (NeuN; white) and oligodendrocytes (MBP; red), as well as (**C**) HSV-1 protein (green), microglia (Iba1; white), and astrocytes (GFAP; red). Nuclei were stained with Hoechst-33342 (blue). Representative images are shown for patient #2 (amygdala/entorhinal cortex; Table 2). Scale bar: 20 μm.

We analyzed whether lytic HSV infection was associated with increased intracellular production of Aβ or pTau, or the deposition of extracellular Aβ plaques in brain sections from HSE patients (Table 1) and, as control for neuronal damage, traumatic brain injury (TBI) (n = 3; Supplementary Table 3) and AD patients (n =3; Supplementary Table 4). TBI-induced neuronal damage is associated with increased production of Aβ, resulting in its accumulation in axonal spheroids as well as plaques in about 30% of patients (37). Indeed, we observed axonal spheroids in all three TBI patients and increased levels of intracellular Aβ protein and (diffuse) extracellular Aβ plaques in two TBI patients (Figure 3A). NFTs and pTau staining were not observed in TBI patients (Figure 3A). Brain sections, containing HSV-infected cells, from two out of five HSE patients did not show Aβ protein/plaques nor pTau protein/NFTs (Table 1). Aβ plaques were observed in one of five HSE patients (patient #2), whereas intracellular accumulations of Aβ protein were detected in three of five HSE patients. NFTs were detected in two of five HSE patients (patient #2 and #5) (Figure 3A and Table 1).

**Figure 3.**
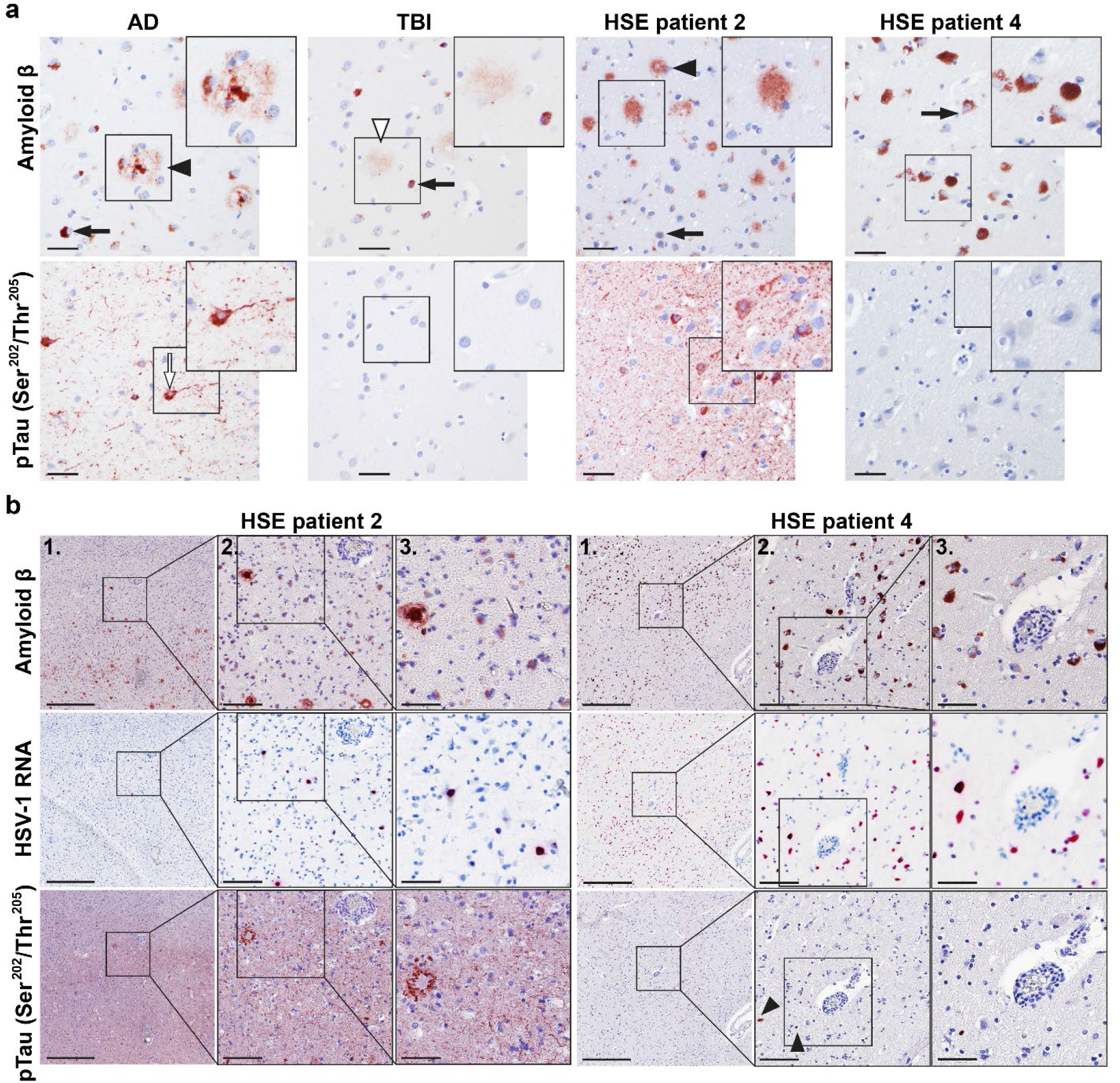
Lytic HSV infection is not consistently associated with aberrant Aβ or pTau expression in brain of herpes simplex virus encephalitis (HSE) patients. (**A**) Brain sections from Alzheimer’s disease (AD; patient 1, Supplementary Table 4), trauma brain injury (TBI; patient 2, Supplementary Table 3) and HSE patients (Table 1) were stained for Aβ and pTau (Ser^202^/Thr^205^) by IHC. Filled and open arrowheads indicate mature senile and diffuse Aβ plaques, respectively. Filled black arrows indicate intracellular Aβ accumulations. Open arrows indicate neurofibrillary tangles. Boxes indicate area shown at higher magnification in the inset. Scale bar: 50 μm. (**B**) Consecutive brain sections from HSE patients were stained for Aβ and pTau (Ser^202^/Thr^205^) protein by IHC and HSV-1 RNA by ISH. Boxes indicate area shown at a higher magnification. Scale bars indicate 500 μm (panels 1), 100 μm (panels 2) and 50 μm (panels 3).

Subsequently, we analyzed the spatial orientation of HSV-infected cells in relation to AD-related pathological changes. Although HSV-infected cells were occasionally observed in close proximity to NFT and/or Aβ plaques, majority of these AD-related pathological changes were not associated with sites of HSV replication in the brains of HSE patient #2 and #5 (Figure 3B). Similarly, intracellular accumulations of Aβ protein were widely spread through brain sections of HSV-1 HSE patients, not restricted to areas with HSV-1 infection and also observed in sections without HSV-infected cells (Figure 3B and Supplementary Figure 2). IF staining was performed to investigate potential co-localization between HSV-1 infected cells and plaques/NFT and to assess intracellular levels of Aβ and pTau in more detail. Again, occasional HSV-infected cells were observed adjacent to Aβ plaques and NFT, but HSV antigen never co-localized with Aβ plaques nor NFT (Figure 4A). In HSE patients #2, #4 and #5 abundant intracellular Aβ protein staining was detected in most HSV-infected neurons, but also in non-infected neurons (Figure 4B). Nuclear pTau staining was observed in some HSV-infected cells in HSE patients containing NFTs (Figure 4C). However, no increased Aβ or pTau levels were observed in HSV-infected neurons in HSE patients #1 and #3 (Figure 4D).

**Figure 4.**
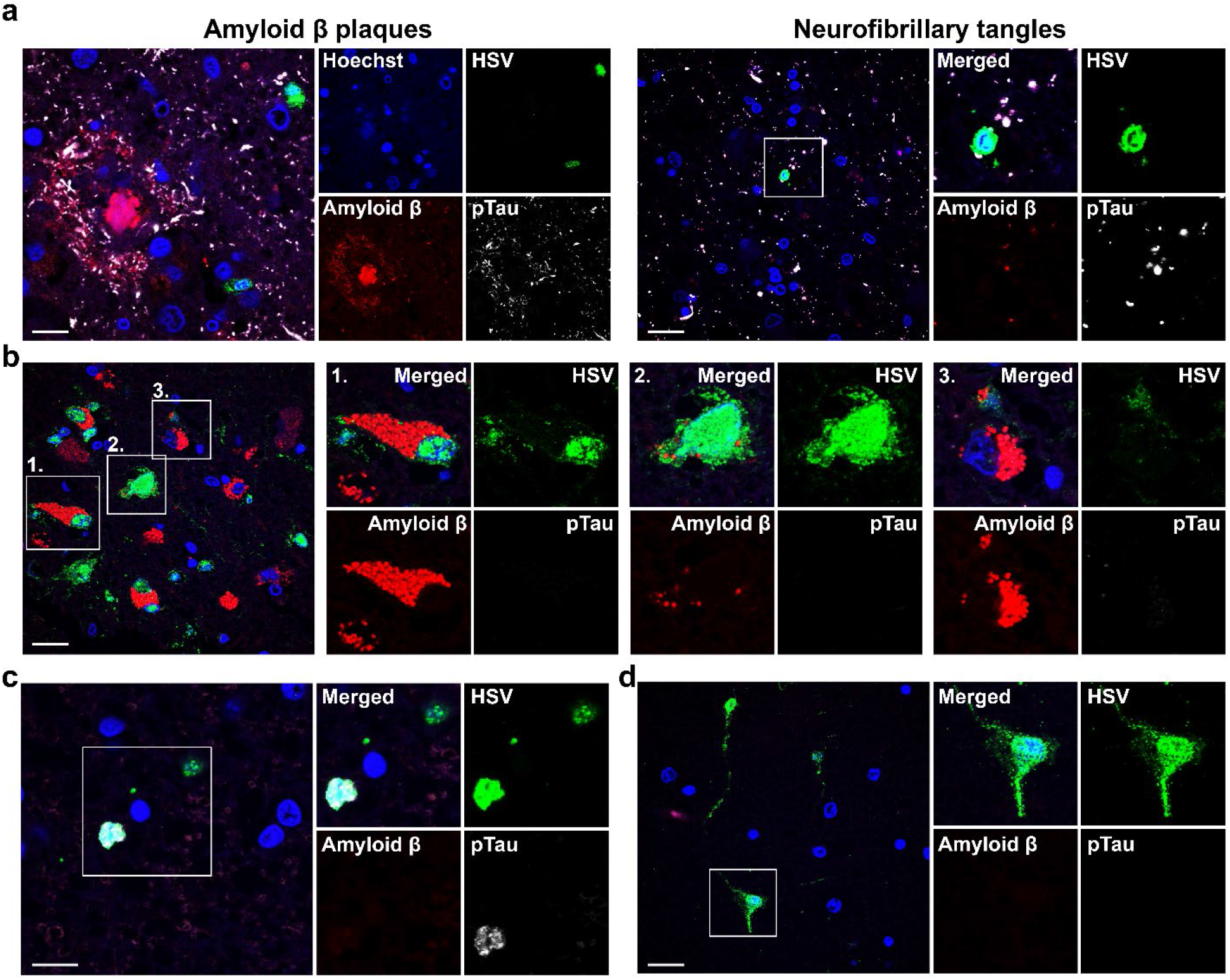
HSV-infected cells do not consistently express aberrant Aβ or pTau in brain of herpes simplex encephalitis (HSE) patients. Brain sections from HSE patients were immunofluorescently stained for HSV protein (green), Aβ (red) and pTau (Ser^202^/Thr^205^; white). Nuclei were stained with Hoechst-33342 (blue). Images are shown for HSE patients #2 (**A** left panel), #5 (**A** right panel, and **C**), #4 (**B**) and #3 (**D**) (Table 1). White boxes indicate areas shown at higher magnification. Scale bars indicate 10 μm (**A**: Amyloid β plaques; and **C**) and 20 μm (**A** right panel, **B** and **D**).

Finally, we investigated whether the widespread increased levels of intracellular Aβ protein, in the absence of diffuse Aβ plaques, in brain of some HSE patients were specific to HSV infection by analyzing brain sections from a case of encephalitis caused by VZV. A 72-year-old, male patient with no known history of dementia presented with VZV encephalitis and died 15 days after onset of disease. Abundant VZV RNA-positive cells were detected (Supplementary Figure 2A), as well as VZV antigen-positive cells, indicative of lytic VZV infection. Aβ plaques and NFT, as well as intracellular Aβ protein were observed in hippocampal sections, whereas only widespread prominent intracellular Aβ protein depositions were observed in medulla oblongata sections (Supplementary Figure 2B). Similar to the HSE brains, we did not observe an association between the spatial orientation of virus-infected cells and AD-related pathological changes (Supplementary Figure 2C). Overall, these results indicate that lytic α-herpesvirus, especially HSV, infection of human CNS neurons *in vivo* is not consistently associated with immediate aberrant Aβ or pTau protein expression.

### Expression of Aβ and pTau in brain of an AD patient with concurrent HSE

To investigate how HSV-1 infection interacts with AD-related pathological changes in the AD brain, we obtained rare brain tissue specimens from an AD patient presenting with HSE. Patient was a 63-year-old female diagnosed with AD who developed HSE and died from septic shock resulting from HSE and aspiration pneumonia 1.5 weeks after hospitalization. Analysis of 6 cortical brain regions revealed intermediate AD-related pathological changes, i.e. amyloid Thal phase 5/5 and Braak NFT stage 4/6 (A3B2), consistent with dementia. HSV antigen-positive cells were abundantly detected in insular and temporal cortical tissue sections and less prominent in parietal and occipital cortex sections (Figure 5A). Importantly, HSV-1 infected cells were often found in close proximity to NFT and especially Aβ plaques (Figure 5B). However, AD-related pathological changes were not different between regions with and without HSV-1 infected cells. HSV-1 infected cells were situated around, but not co-localized with, Aβ plaques and NFTs (Figure 5C). Similar to HSE patients, increased levels of intracellular Aβ protein were present in most HSV-infected cells, as well as non-infected cells (Figure 5D). Thus, AD and HSV infection may affect the same brain regions, but we did not observe enhanced AD-related pathology in relation to HSV infection.

**Figure 5.**
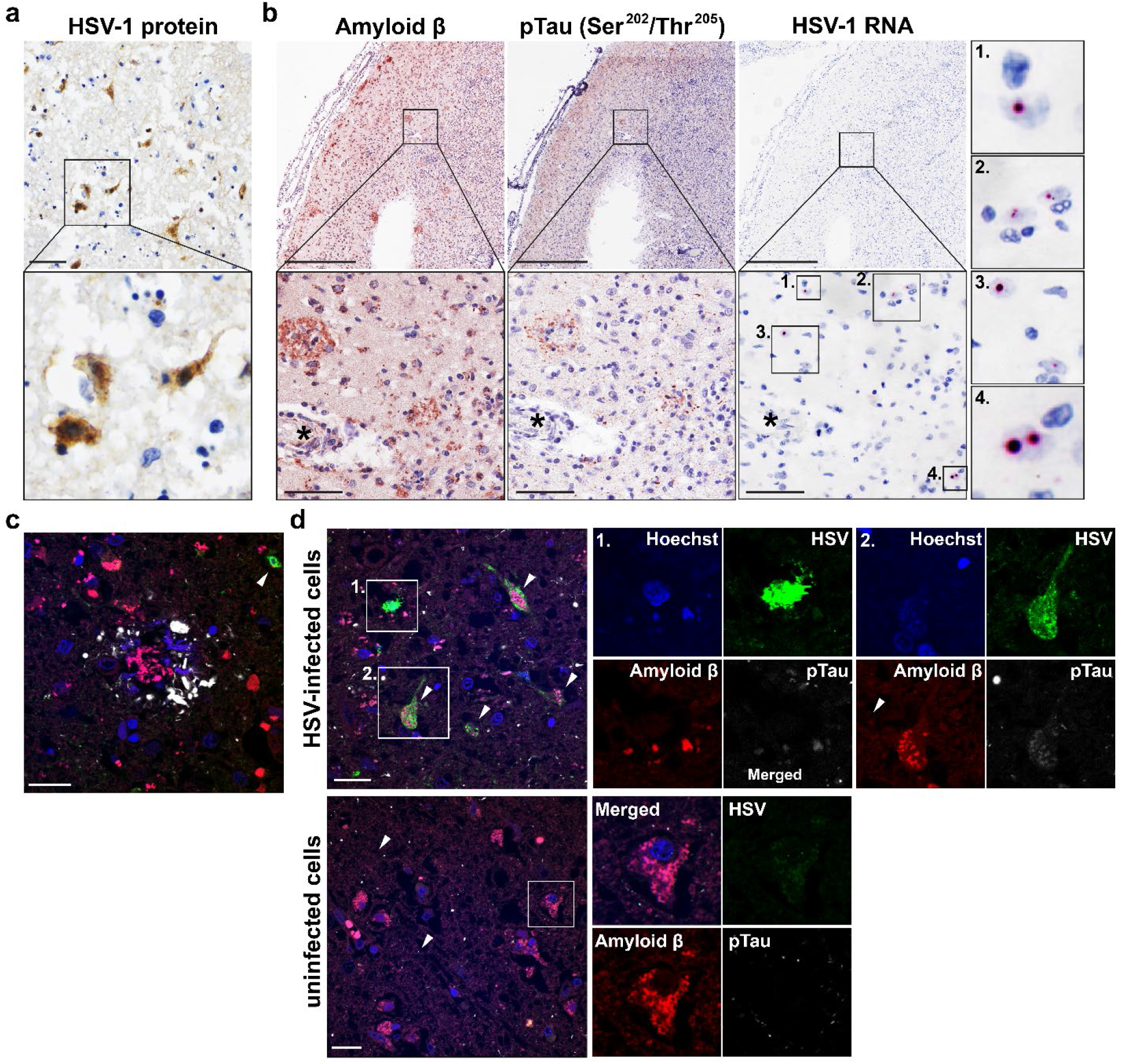
Detection of HSV-infected cells in close proximity to Aβ plaques in the brain of an Alzheimer’s disease patient with herpes simplex encephalitis. **A** Brain section stained for HSV protein (brown) by immunohistochemistry (IHC). Scale bar indicates 50 μm. **B** Consecutive brain sections were stained for Aβ (4G8) and pTau (Ser^202^, Thr^205^) by IHC or stained for HSV-1 RNA by in situ hybridization (ISH). Boxes indicate areas shown at higher magnification. Asterisks indicate same blood vessel in consecutive sections. Scale bars indicate 500 μm (low magnification) and 100 μm (high magnification). **C-D** Brain sections were immunofluorescently stained for HSV protein (green), Aβ (red) and pTau (Ser^202^/Thr^205^; white) protein. Nuclei were stained with Hoechst-33342 (blue). Scale bar: 20 μm.

### Neuroinflammation in HSE patients with and without AD-related pathology

Because neuroinflammation is a major contributor to neurodegeneration in AD (38), we hypothesized that inflammatory cells rather than HSV replication itself could be associated with specific AD-related pathological changes. Brain tissue sections of all HSE patients, including the AD patient with HSE, indeed demonstrated widespread inflammation and gliosis in comparison to AD patients without viral encephalitis (Figure 6A and Supplementary Table 4). HSE patients and the AD patient with concurrent HSE showed prominent perivascular cuffs, mainly composed of CD45^POS^ CD3^NEG^ Iba1^NEG^ mononuclear cells – most likely macrophages or histiocytes – and lower numbers of CD45^POS^ CD3^POS^ T-cells (Figure 6B). Astrogliosis was observed in the brain parenchyma of all HSE patients, irrespective of Aβ or pTau expression pattern. This was also observed in the AD patient with HSE, but not as extensively in the AD patients (Figure 6C). Notably, while astrocytes interacted with Aβ plaques of all AD patients and HSE patient #2, more prominent GFAP staining was observed in astrocytes embracing Aβ plaques in the AD patient with HSE (Supplementary Figure 4). Additionally, extensive microgliosis was observed in the brain parenchyma of the HSE patients and AD patient with HSE, but not AD patients (Figure 6D). Both amoeboid (Figure 6D) and ramified microglia (Supplementary Figure 5) were observed throughout the tissue sections and in association with Aβ plaques and/or pTau in the AD patient with concurrent HSE and the HSE patients #2 and #5 (Table 2). Microglia were mostly amoeboid in HSE patients #1 and #3, whereas predominantly ramified microglia were present in HSE patient #4 (Figure 6D). Thus, although lytic HSV infection of the CNS induced significant neuroinflammation, the overall pattern of inflammation and gliosis was not associated with specific Aβ or pTau phenotypes.

**Figure 6.**
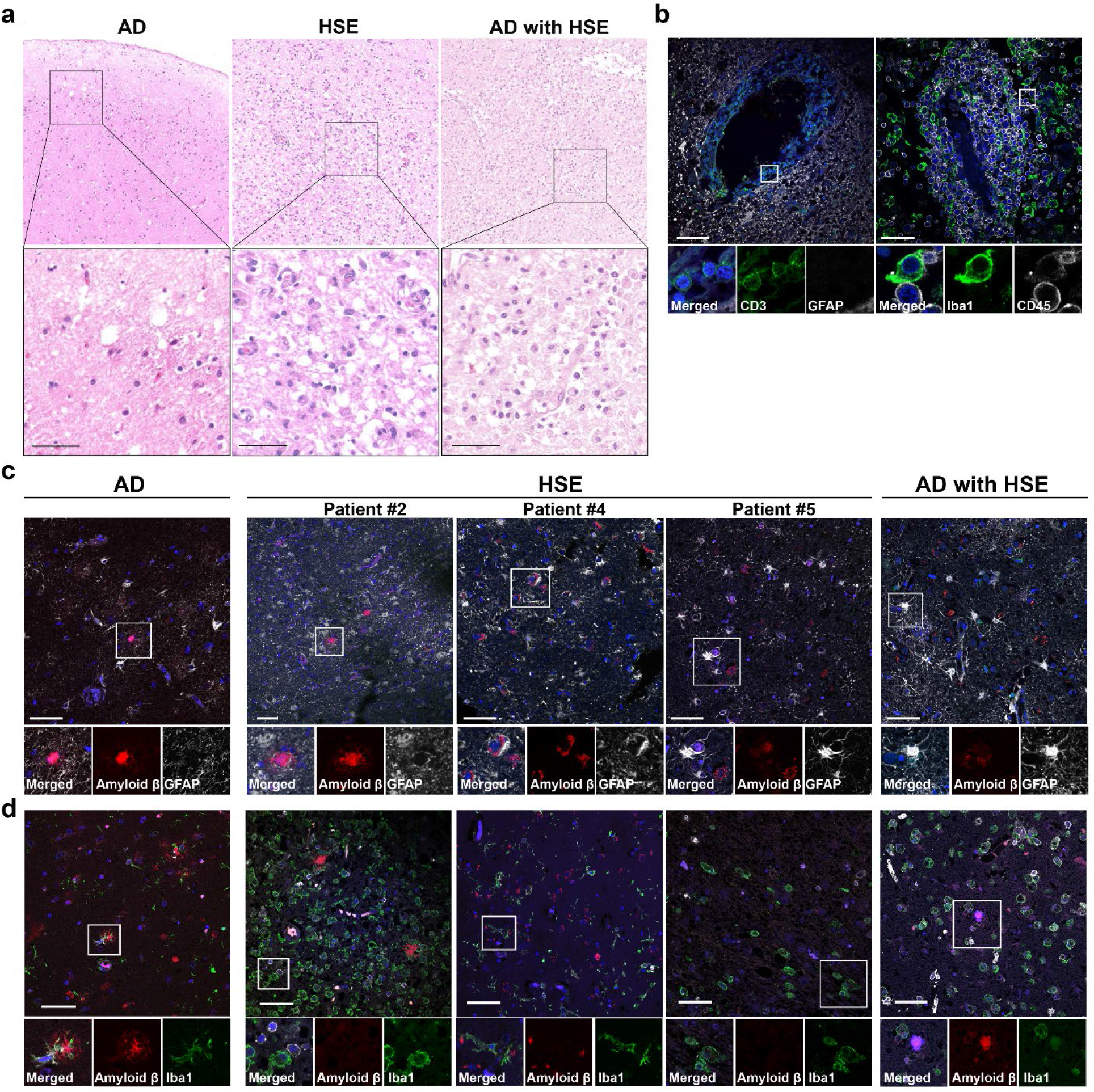
Neuroinflammation is similar in herpes simplex encephalitis (HSE) patients with and without Aβ plaques and/or NFT. Brain sections from Alzheimer’s disease (AD) patients (n=3, Supplementary Table 4), HSE patients (n=4, Table 1) and an AD with HSE patient (n=1) were stained with (**A**) hematoxylin and eosin staining or IF stained for: (**B**) CD3 (green) and GFAP (white) (left panel) or Iba1 (green) and CD45 (white) (right panel); (**C**) Aβ (red) and GFAP (white); or (**D**) Aβ (red), Iba1 (green) and CD45 (white). Nuclei were stained with Hoechst-33342 (blue). Images are shown for the AD patient with HSE (**B** – **D**), AD patient #1 (**C**, **D**; Supplementary Table 4) and indicated HSE patients. Boxes indicate areas shown at higher magnification. Scale bar: 50 μm.

## DISCUSSION

Cumulative evidence suggests that the endemic human neurotropic virus, HSV-1, could play a role in the pathogenesis of AD (14,15,18–21,39). Here, we used clinical specimens from HSV-infected individuals to investigate the relationship between HSV infection and Aβ and pTau expression within human neurons *in vivo*. We report that latent HSV infection is not associated with Aβ or pTau expression in TG neurons located in the PNS. Similarly, lytic HSV infection in the CNS was not consistently associated with increased expression of intracellular Aβ or pTau proteins nor with the deposition of Aβ plaques. Analysis of unique material from an AD patient with concurrent HSE showed that the same brain regions are often affected by HSV and AD-related pathology, but did not reveal exacerbated Aβ or pTau production in relation to regions containing virus-infected cells.

Previous studies show that increased HSV IgM plasma levels and possibly anti-HSV IgG avidity can be a measurement of HSV reactivation that correlates with AD development (26,40). One potential explanation is that genetic risk factors associated with AD development, especially *APOE4* and paired immunoglobulin-like type 2 receptor alpha (*PILRA*), directly impact lytic HSV-1 infection. The *APOE4* allele is associated with increased frequency of HSV-1 DNA detection in brain of AD patients and symptomatic HSV-1 reactivation (cold sores) (11,41). By contrast, asymptomatic oral HSV-1 shedding was not affected by *APOE* genotype (41), suggesting that *APOE4* does not directly influence virus reactivation but may play a role in peripheral control of HSV infection. In this study, we did not observe differences in latent HSV-1 DNA load in the TG of AD patients compared to controls or in *APOE4* carriers compared to non-carriers. Collectively, these data indicate that HSV-1 reactivation, rather than HSV-1 infection, is a risk factor for AD development, especially in *APOE4* carriers.

Detailed analysis of rare human HSE brain specimens demonstrated that lytic HSV infection was not associated with increased levels of intraneuronal Aβ or Aβ plaque deposition in human neurons *in vivo*. Although we observed accumulation of intraneuronal Aβ protein in three HSE and one VZV encephalitis patient, Aβ depositions were widely present in both virus-infected and non-infected neurons and tissue sections. These findings are consistent with the ubiquitous and progressive accumulation of intraneuronal Aβ in the absence of extracellular Aβ plaques and NFT that is observed with aging (42). Similarly, we only observed Aβ plaques in one HSE patient and locally (hippocampus, but not medulla oblongata) in one VZV encephalitis patient. The elderly age of these two patients, lack of an association between sites of virus replication and Aβ plaques, and the presence of both senile plaques and NFT suggests that these pathological changes were most likely part of an ongoing development of AD, rather than induced by HSV infection.

HSV particles bind to Aβ protein and catalyze Aβ42 oligomerization, leading to Aβ aggregates that physically entrap virus particles (7,43). Consistent with the idea of HSV initiating the deposition of Aβ plaques in the CNS, a previous study used in situ PCR to detect HSV-1 DNA in the brain of AD patients and controls to demonstrate that HSV-1 preferentially co-localizes with Aβ plaques in AD patients (12). Here, we did not detect viral antigen nor RNA within Aβ plaques in brain of elderly individuals with HSV-1 or VZV encephalitis. Although HSV-infected cells were often found in close proximity to Aβ plaques and/or NFT in the brain of the AD patient with HSE, AD-related pathology was not restricted to regions with HSV-infected cells and also in this patient we did not observe HSV antigen or RNA in Aβ plaques. HSV-1 infection rapidly induces Aβ plaque deposition in the brain of genetically susceptible AD mouse models, but repeated HSV-1 reactivation and lytic virus replication in brain results in the progressive accumulation of both senile plaques and NFT (21). While HSE is extremely rare in humans, asymptomatic HSV reactivation and viral spread to the CNS, as measured by the presence of viral DNA, occurs frequently (11,12). Our data indicate that lytic HSV infection does not directly induce the formation of Aβ plaques in the human brain, possibly due to the extensive neuronal cell death observed in HSE patients. Instead, repeated exposure to abortive HSV infections may be required to induce persistent Aβ plaques or could enhance Aβ oligomerization in existing plaques.

HSV infection of diverse human neuron models *in vitro* leads to the hyperphosphorylation of tau protein, including Ser^202^/Thr^205^, by cellular cyclin-dependent kinases and relocation of pau to the nucleus (18–20). We report that HSV-infected neurons in brain of human HSE patients occasionally expressed nuclear pTau, but only in individuals presenting with NFT. However, NFTs were not restricted to areas containing HSV-infected cells in HSE patients and the AD patient with HSE. These findings are in agreement with prior studies in 3×Tg-AD mice (prone to develop NFT (44)), in which the degree and residues involved in tau phosphorylation in response to HSV infection varied between brain regions, and progressively increased with repeated viral reactivation events (21). Possibly, the heterogeneity of neurons *in vivo* – including the expression of specific cyclin-dependent kinases – determines their differential susceptibility to HSV-induced aberrant tau phosphorylation and cellular localization.

Specific microglia and astrocyte populations colocalize with Aβ plaques in brain of AD patients (45–47) and are thought to be involved in both early and advanced stages of AD pathogenesis (48,49). Lytic HSV infection induces robust innate immune responses in brain, involving both microglia and astrocytes, which control ongoing virus replication, but also, cause immunopathology (50). Prominent microgliosis and astrogliosis were present in all HSE patients, but we did not observe overall differences in glia cell morphology or abundance in patients with or without Aβ plaques and/or NFT. Interestingly, GFAP staining tended to be more abundant in Aβ plaque-associated astrocytes in brain sections from the patient with combined AD and HSE, compared to patients with either AD or HSE (Supplementary Figure 4). Although these observations suggest that HSV infection may influence reactive astrocyte function in brain of AD patients, more detailed analyses in more patients or experimental animal models are warranted.

In conclusion, we demonstrate that latent and lytic HSV infection of human neurons *in vivo* is not consistently associated with aberrant Aβ or pTau expression. Importantly, our results do not challenge the increasing epidemiological and functional evidence that implicates a role of HSV-1 infection in the pathogenesis of AD. Human *in vitro* neuron cultures and murine AD models highlight potential mechanisms by which HSV infection can contribute to the initiation, perpetuation, or deterioration of AD. However, our data suggest that the human CNS could be more resilient to HSV-induced AD-related neuropathology than previously anticipated. Future studies comparing the effects of HSV infection on the healthy ageing brain and AD brain may provide valuable insight into mechanisms by which HSV may affect AD pathogenesis.

## ACKNOWLEDGMENTS

We thank Dr. Georges Verjans for critical discussion of the data and Tamana Khemai-Mehraban for technical assistance. Research reported in this publication was in part supported by the National Institute of Allergy and Infectious Diseases of the National Institutes of Health under Award Number R01AI151290 (W.J.D.O.). The content is solely the responsibility of the authors and does not necessarily represent the official views of the National Institutes of Health. This study was funded in part by a Human Disease Model Award 2020 (Erasmus MC).

## SUPPLEMENTARY TABLES AND SUPPLEMENTARY FIGURE LEGENDS

**Supplementary Table 1.**
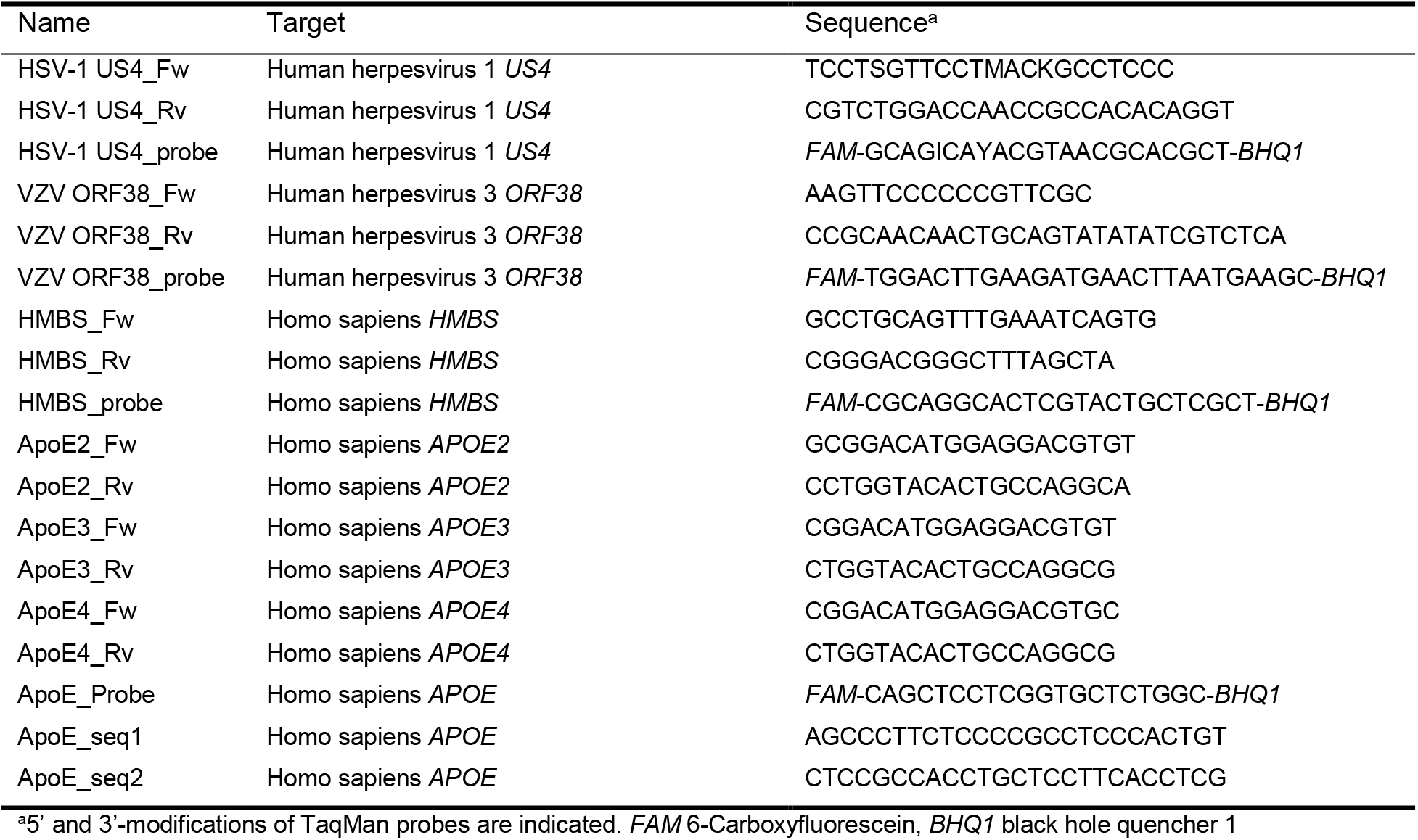
Primers and probes used in this study

**Supplementary Table 2.** FFPE TG samples used in this study

| Subject | Age<sup>a</sup> | Gender<sup>b</sup> | PMI<sup>c</sup> | Cause of death | Neurological disease | HSV-1 status<sup>d</sup> |
| --- | --- | --- | --- | --- | --- | --- |
| 1 | 81 | F | 3:35 | Cachexia and dehydration | Alzheimer's disease | negative |
| 2 | 82 | F | 5:55 | Cardiac arrest | Alzheimer's disease | negative |
| 3 | 89 | F | 4:30 | Peritonitis | Alzheimer's disease | positive |
| 4 | 87 | F | 8:10 | Uremia by dehydration | Alzheimer's disease | positive |
| 5 | 86 | F | 5:15 | Cachexia and dehydration | Alzheimer's disease | positive |
| 6 | 81 | M | 8:21 | Euthanasia | Parkinson's disease | positive |
| 7 | 62 | F | 12:35 | Cachexia with pulmonary<br>insufficiency | Multiple sclerosis | positive |
| 8 | 80 | F | 9:30 | Cardiac arrest | Bipolar disorder | positive |
| 9 | 103 | F | 6:35 | Dehydration and pneumonia | Tauopathy | positive |
| 10 | 89 | M | 6:50 | Urosepsis | non-demented control | positive |

<sup>a</sup>Age in years; <sup>b</sup>F, female; M, male; <sup>c</sup>PMI, post-mortem interval; <sup>d</sup>HSV-1 infection status based on serology and HSV-1 LAT ISH

**Supplementary Table 3.**
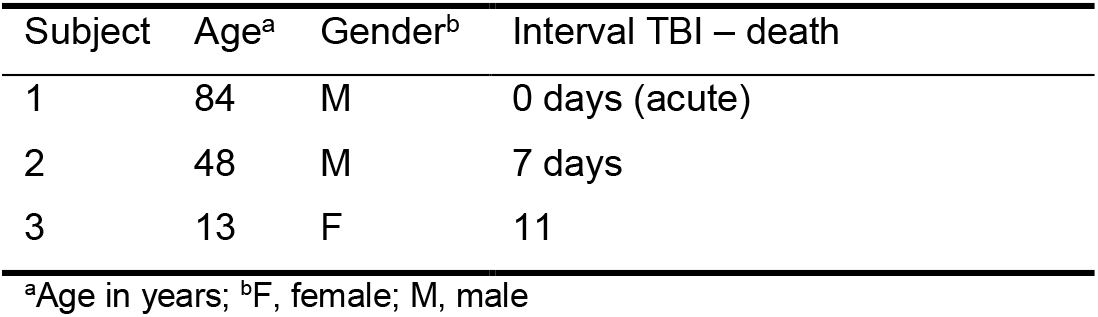
TBI patients used in this study

| Subject | Age<sup>a</sup> | Gender<sup>b</sup> | Interval TBI – death |
| --- | --- | --- | --- |
| 1 | 84 | M | 0 days (acute) |
| 2 | 48 | M | 7 days |
| 3 | 13 | F | 11 |

<sup>a</sup>Age in years; <sup>b</sup>F, female; M, male

**Supplementary Table 4.** FFPE AD samples used in this study

| Subject | Age<sup>a</sup> | Gender<sup>b</sup> | PMI<sup>c</sup> | Cause of death | Neurological disease | HSV status<sup>d</sup> |
| --- | --- | --- | --- | --- | --- | --- |
| 1 | 88 | M | 4:40 | Cachexia and dehydration | Alzheimer's disease | positive |
| 2 | 74 | M | 8:00 | Dehydration by possible<br>cystitis | Alzheimer's disease | positive |
| 3 | 86 | M | 05:10 | Cachexia and dehydration | Alzheimer's disease | positive |

<sup>a</sup>Age in years; <sup>b</sup>F, female; M, male; <sup>c</sup>PMI, post-mortem interval; <sup>d</sup>HSV infection status based on HSV-specific IgG serology data

---

**Supplementary Fig. 1.**
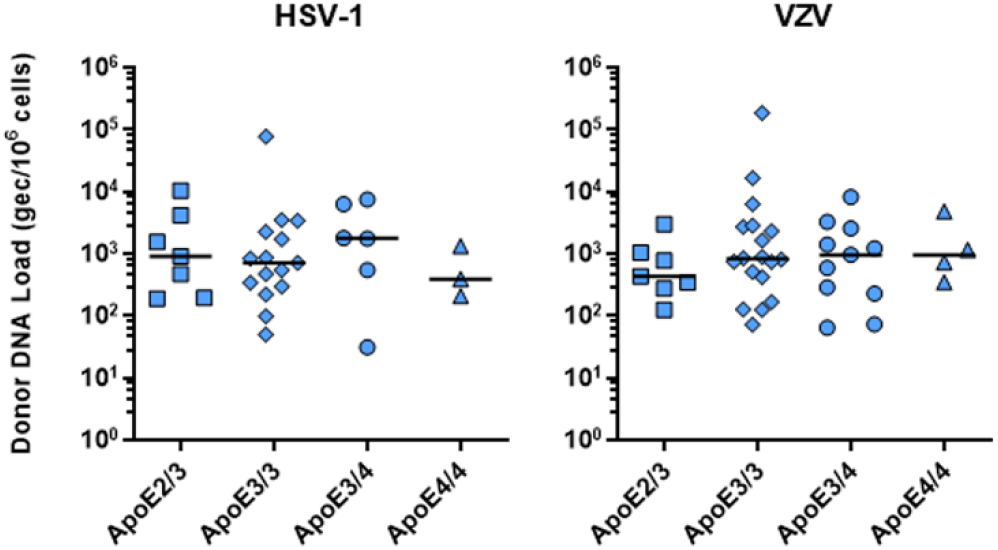
Quantification of latent HSV-1 and VZV DNA load in human TG stratified on *APOE* genotype. HSV-1- and VZV-specific qPCR and *APOE* genotyping was performed on DNA extracted from the trigeminal ganglia (TG) of AD patients and controls.

**Supplementary Fig. 2.**
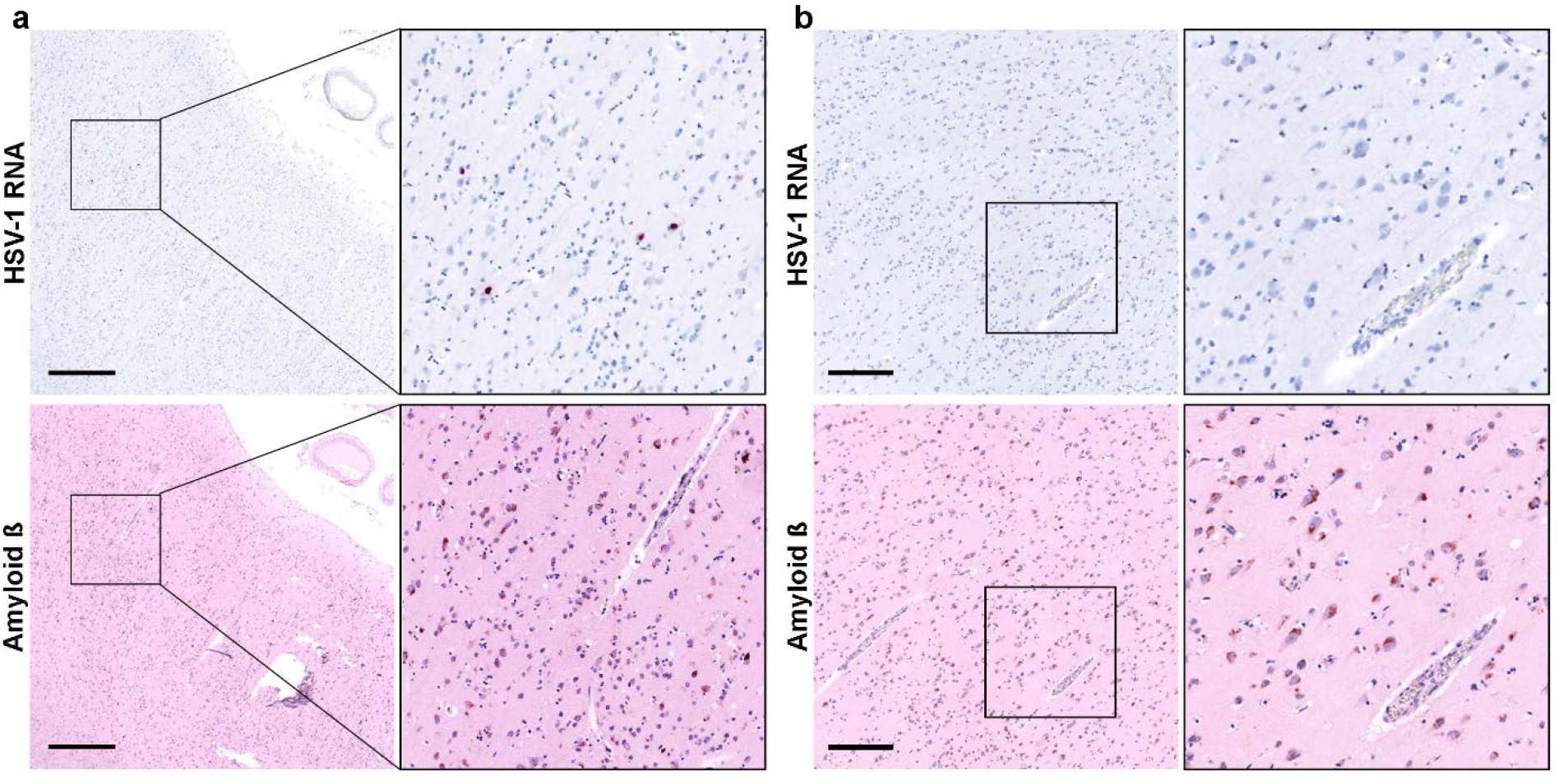
Intraneuronal accumulation of Aβ protein is not restricted to areas of HSV-1 infection in brains of HSE patients. Consecutive slides were stained for HSV-1 RNA by ISH and Aβ by IHC. Aβ can be seen in areas with HSV-1 RNA (**a**) and in areas without HSV-1 RNA (**b**). Data shown for HSE donor 5. Scale bar: 500 μm (**a**) and 250 μm (**b**).

**Supplementary Fig. 3.**
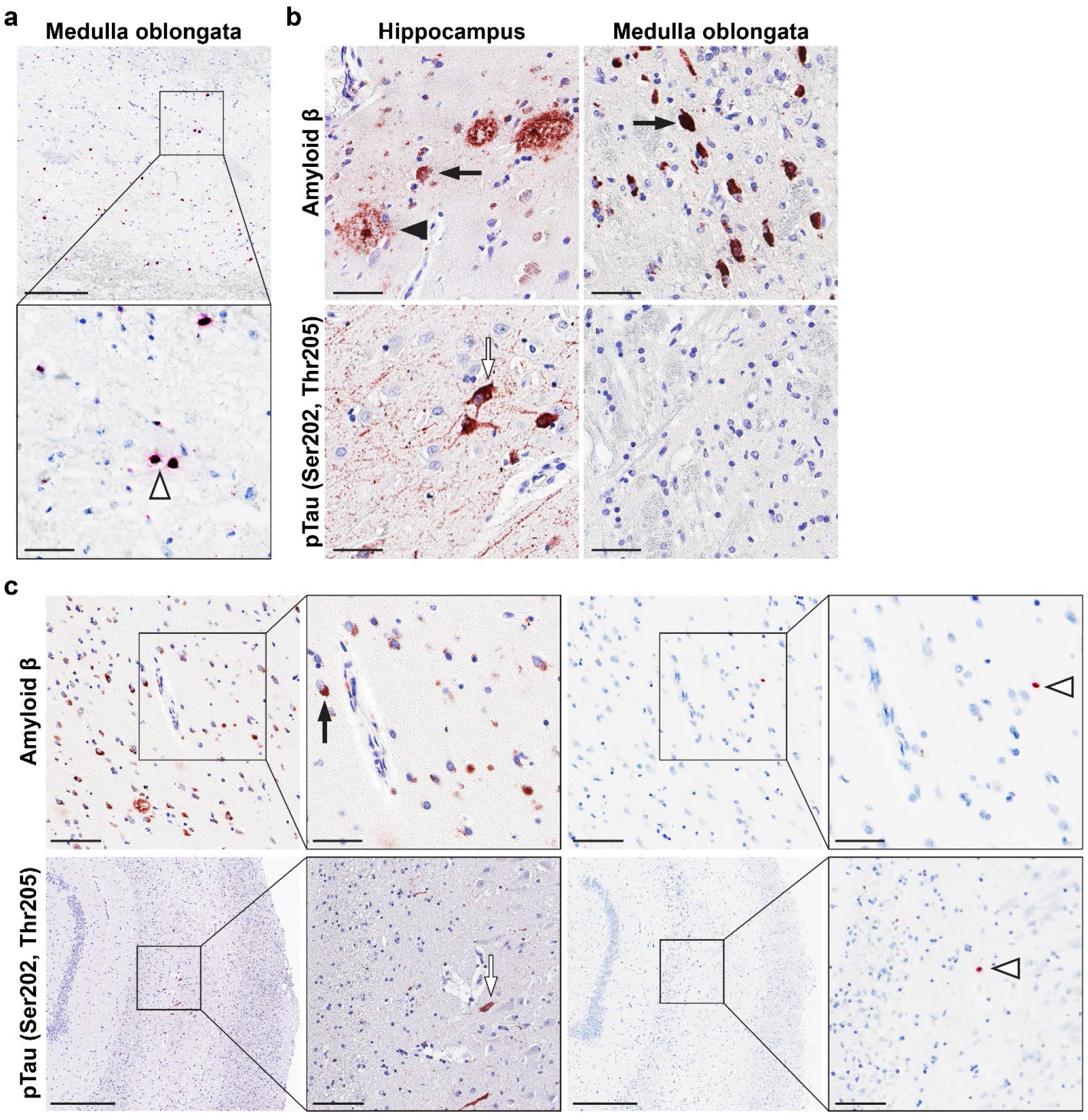
Lytic VZV infection is not associated with Aβ plaques or NFT in brain of a VZV encephalitis patient. **a** Brain section stained for VZV RNA by ISH. Box indicates area shown at higher magnification. Open arrowhead indicates examples of VZV RNA-expressing cells. Scale bars indicate 250 μm (top) and 50 μm (bottom). **b** Brain sections IHC stained for Aβ and pTau (Ser^202^/Thr^205^). Scale bars indicate 50 μm. **c** Consecutive brain sections were IHC stained for Aβ and pTau (Ser^202^/Thr^205^) or stained for HSV-1 RNA by ISH. Boxes indicate areas shown at higher magnification. Filled arrow indicates intracellular Aβ protein, open arrow indicates NFT and open arrowhead indicates VZV RNA-expressing cells. Scale bars indicate 50 μm (Aβ, high magnification), 100 μm (Aβ, low magnification; pTau high magnification) and 500 μm (pTau, low magnification).

**Supplementary Fig. 4.**
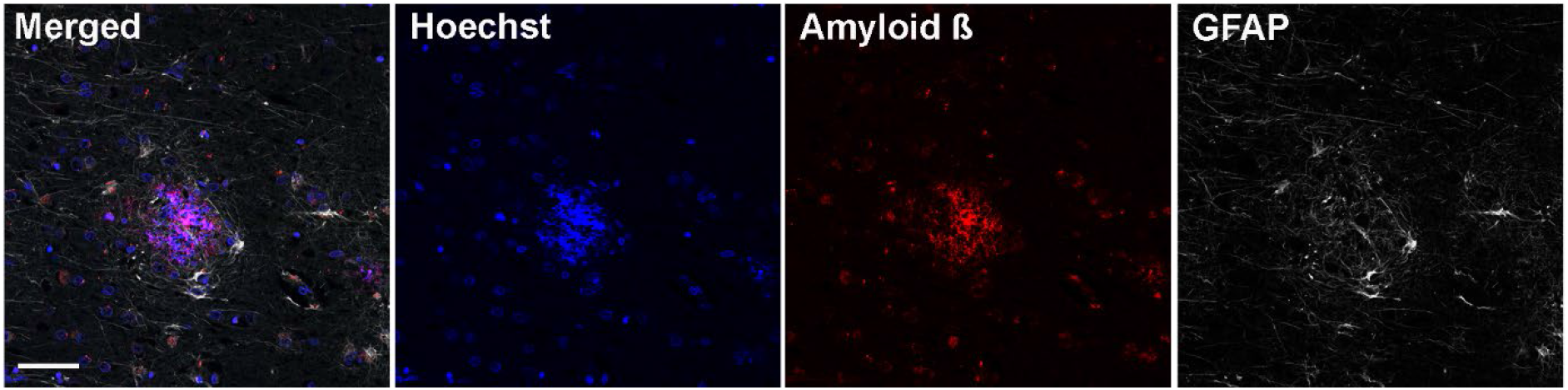
Prominent GFAP staining of astrocytes interacting with Aβ plaques brain of an AD patient with concurrent HSE. IF staining for Aβ (red), GFAP (white) and nuclei (Hoechst-33342; blue). Scale bar: 50 μm.

**Supplementary Fig. 5.**
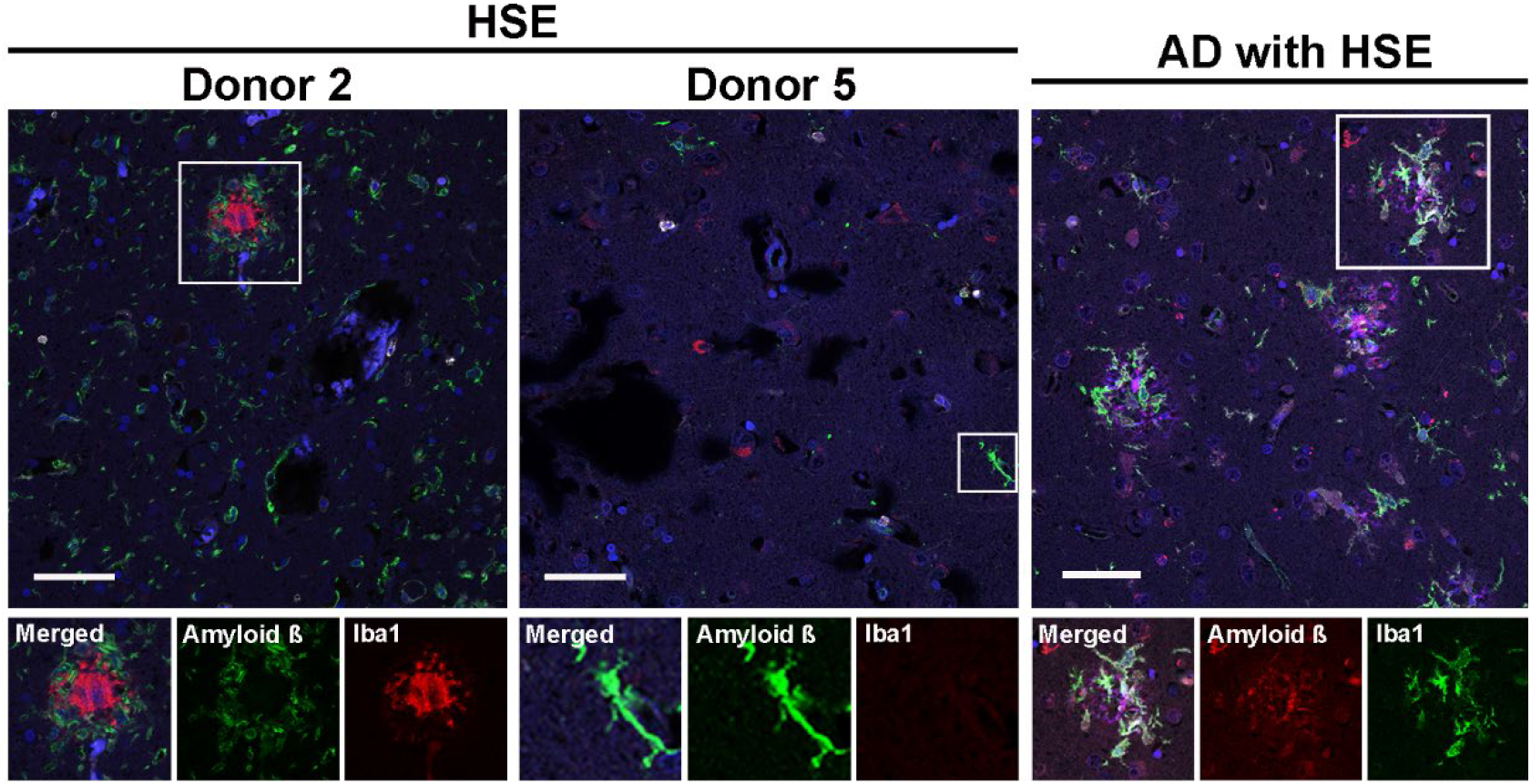
Microglia morphology and density in the brain of HSE patients and AD patient with HSE. Brain tissue sections were IF stained for Aβ (red), Iba1 (green) and nuclei (Hoechst-33342; blue). Boxes indicate areas shown at higher magnification. Scale bar: 50 μm.

